# Machine Learning–Informed Predictions of Nanoparticle Mobility and Fate in the Mucus Barrier

**DOI:** 10.1101/2022.03.04.483046

**Authors:** Logan Kaler, Katherine Joyner, Gregg A. Duncan

**Affiliations:** Biophysics Program, University of Maryland, College Park, MD 20742, USA; Fischell Department of Bioengineering, University of Maryland, College Park, MD 20742, USA

## Abstract

Diffusion and transport of nanomaterials through mucus, is of critical importance to many basic and applied areas of research such as drug delivery and infectious disease. However, it is often challenging to interpret the dynamics of nanoparticles within the mucus gel due to its inherently heterogeneous microstructure and biochemistry. In this study, we measured the diffusion of densely PEGylated nanoparticles (NP) in human airway mucus *ex vivo* using multiple particle tracking and utilized machine learning to classify NP movement as either traditional Brownian motion (BM) or one of two different models of anomalous diffusion, fractional Brownian motion (FBM) and continuous time random walk (CTRW). Specifically, we employed a physics-based neural network model to predict the modes of diffusion experienced by individual NP in human airway mucus. We observed rapidly diffusing NP primarily exhibit BM whereas CTRW and FBM exhibited lower diffusion rates. Given the use of muco-inert nanoparticles, the observed transition from diffusive (BM) to sub-diffusive (CTRW/FBM) motion is likely a result of patient-to-patient variation in mucus network pore size. Using mathematic models that account for the mode of NP diffusion, we predicted the percentage of nanoparticles that would cross the mucus barrier over time in human airway mucus with varied total solids concentration. We also applied this approach to explore the transport modes and predicted fate of influenza A virus within human mucus. These results provide new tools to evaluate the extent of synthetic and viral nanoparticle penetration through mucus in the lung and other tissues.

## INTRODUCTION

Airway mucus acts as a natural filter to capture inhaled particulates by coating lung epithelial surfaces and is a critical component of innate defenses in the respiratory tract. Mucus is comprised of secreted gel-forming mucins which form into a biopolymer network with low viscous and elastic moduli allowing for efficient transport at airway surfaces (Song et al., 2020). The mucus layer is continually transported by the coordinated beating of cilia on airway epithelial cells, which is the primary mechanism of removing mucus from the lungs, referred to as mucociliary clearance (MCC) (Huck et al., 2022). The network structure of the mucus gel enables the trapping and removal of micro– and nanoscale particles via MCC. MCC provides our body’s first line of defense against inhaled pathogens, such as respiratory viruses (Fahy and Dickey, 2010). Similarly, MCC may also limit the bioavailability of therapeutic nanocarriers posing a significant challenge in inhaled drug delivery applications (Ernst et al., 2017)

Whether therapeutics or pathogens, the ability for particles to overcome the mucus barrier is dependent on the particle’s surface chemistry, size, shape, and rigidity. Particles with a positively charged or hydrophobic surface adhere strongly to the mucus network due to net negatively charged and hydrophobic domains within mucins (Song et al., 2020). It has been shown in previous work that mucoadhesion can be limited by coating hydrophobic and/or charged nanoparticles in a layer of polyethylene glycol (PEG) (Schneider et al., 2017). PEG is hydrophilic, net-neutral polymer which prevents hydrophobic and charge-mediated interactions with mucins, allowing particles to move more freely through the mucus layer. A previous study has also shown that nanoparticles with peptide coatings having a net-neutrally charged amino acid sequence are capable of rapid diffusion through the mucus network (Li et al., 2013). For pathogens like influenza, specific interactions of viral particles with mucin glycans can potentially lead to their entrapment in the mucus layer (Iverson et al., 2020). Depending on their dimensions, nanoscale particles may also be physically immobilized within the mucus gel when much larger than the mucus pore size which ranges from 100-500 nm (Witten and Ribbeck, 2017). Particles that are smaller than the mesh network are likely to diffuse at a rate that corresponds to the viscosity of the fluid–filled pores between the network fibers (Witten et al., 2018). However, heterogeneity of the mucus network can also lead to non-uniformity in nanoparticle diffusion with more rapid movement in regions with decreased mucin density whereas regions with increased mucin density can significantly hinder and immobilize particles even when densely coated in PEG (Huck et al., 2022).

To characterize the diffusion and transport of nanomaterials, multiple particle tracking (MPT) analysis is a commonly used technique to directly measure the diffusion rate of individual nanoparticles within mucus and other complex biological fluids (Schuster et al., 2015). However as a result of the heterogeneity in the mucus gel previously noted, the type of diffusion may vary substantially as of function of spatial position. This complicates interpretation of MPT experiments as the ability of nanoscale particles to navigate through the mucus barrier will be highly dependent on the mode of diffusion. Furthermore, MPT analysis is most often limited to timescales on the order of seconds whereas nanomaterial diffusion and transport through mucus in the lung occurs over physiological timespans over minutes to hours (Duncan et al., 2016a). Thus, new analytical pipelines have been sought to connect MPT analysis to physical models of traversal time through the mucus barrier (Newby et al., 2018). However, these models often require generalized assumptions about the mode of diffusion which are unlikely to fully capture NP dynamics within the highly complex mucus microenvironment.

Towards this end, we combine machine learning and mathematical models to predict the passage times of nanoparticles and viruses across the mucus barrier. Specifically, we use a previously developed machine learning-based analysis called MotionNet (MoNet) to classify the type of diffusion exhibited in individual particle trajectories (Jamali et al., 2021). With these classifications determined, the percentage of particles able to penetrate a mucus layer with physiological thickness of 10 μm are calculated using diffusion mode-dependent analytical expressions (Erickson et al., 2015). We applied this analytical approach to interpret trajectories of PEGylated fluorescent nanoparticles (NP) in human mucus acquired using MPT in 30 distinct human mucus samples. In addition, we evaluated expected passage times for influenza in human mucus using our recently published dataset (Kaler et al., 2022). The results of this work provide an approach to predict the timescales which synthetic and viral nanoparticle diffuse through the mucus layer in the airway and other mucosal tissues.

## RESULTS

### Predicted classification of muco-inert nanoparticles in human mucus samples

We utilized the MoNet analysis developed in previous work (Jamali et al., 2021) on experimental MPT data of 100 nm NP diffusion within human mucus samples collected from endotracheal tubes. Using this analysis, we considered three diffusion modes: Brownian motion (BM), fractional Brownian motion (FBM), and continuous time random walk (CTRW). BM is a standard model for diffusion in a Newtonian fluid (e.g. water) where a particle undergoes a random walk, taking a step left or right with equal probability each time it moves (Erickson et al., 2015). BM would most likely be reflective of free diffusion within aqueous regions of the gel. FBM and CTRW are both sub-diffusive models but driven through distinct processes. CTRW motion is characterized by random jumps in time and space leading to “hopping” NP diffusion. In FBM, NP follow a random walk but subsequent steps are anti-correlated, meaning that there is a higher probability that the next step will be in the opposite direction than the previous step (Jamali et al., 2021). While FBM is more commonly observed in viscoelastic fluids (Valentine et al., 2001), “hopping” motion has been more recently considered as a mode of NP diffusion through biopolymer matrices (Cai et al., 2015). Based on this analysis, we observed all 3 forms of diffusion in our dataset with representative trajectories for each type of motion shown in **Figure 1A**. A representative walkthrough of the MoNet analysis is shown in **Figure 1B-C** for data collected in an individual patient sample. The MoNet analysis determines the probability that an individual particle will exhibit one of the three types of motion (**Figure 1B**) and then predicts the type of motion for that individual particle, based on these probabilities (**Figure 1C**). Using this analysis, we found the predominant type of NP motion in all samples tested to be FBM, with a smaller fraction of NP exhibiting CTRW or BM (**Figure 1D**).

**Figure 1.**
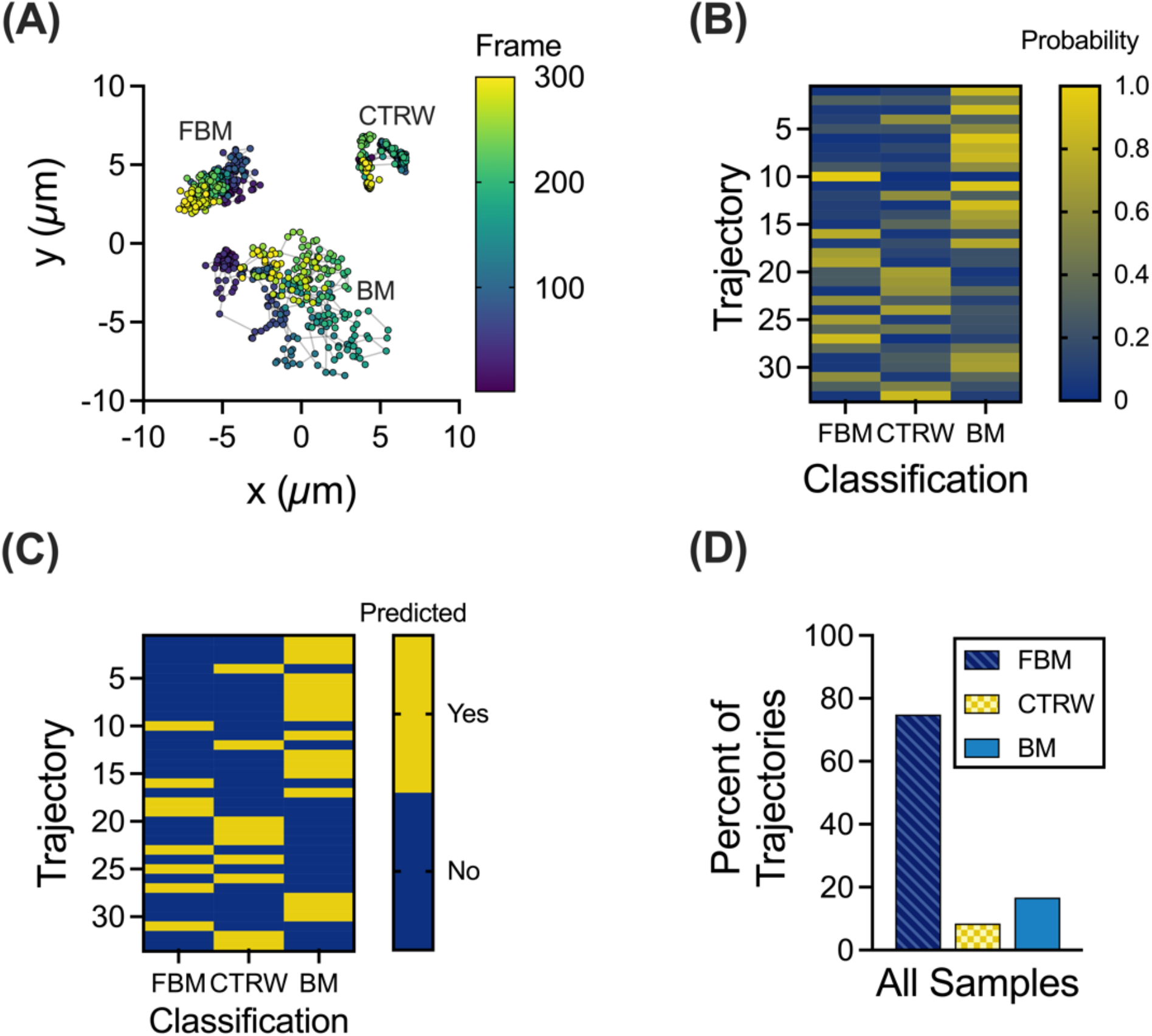
Machine learning based analysis to classify NP trajectories. **(A)** Representative trajectories for each NP motion type; FBM, BM, and CTRW. Trajectory color corresponds to frame of video with purple as the first frame and yellow as the last frame. Nanoparticle motion was captured at a frame rate of 33.3 frames per second. (**B-C**) Representative walk-through of MoNet analysis, with the probabilities for each trajectory **(B)**, and the predicted motion **(C)**. **(D)** Percent of trajectories exhibiting FBM, CTRW, or BM across all samples (n = 30 samples).

### Classification of trajectories correlates with mucus network size

Due to inherent sample-to-sample variability in mucus properties (Duncan et al., 2016b; Joyner and Duncan, 2019; Markovetz et al., 2019), we compared the type of NP diffusion exhibited in individual patients. Based on measured logarithm based 10 (log_10_) of the mean squared displacement (MSD) values at a lag time of 1 second (log_10_[MSD_1s_]), a measure of diffusivity (**Figure 2A**), we found that the diffusivity varied dramatically from sample to sample with median log_10_[MSD_1s_] spanning over ~3 orders of magnitude. We utilized the MoNet analysis on the individual trajectories of each sample and compared the percent of NP exhibiting each type of motion with the estimated pore size (*ξ*) for each sample (**Figure 2B**). Interestingly, we observed that as the pore size in individual patient samples decreased the percent of FBM NP increased and the percent of BM NP decreased. CTRW diffusion was observed the least frequently in individual samples (0–33.3%) but was observed in 20 out of 30 samples tested.

**Figure 2.**
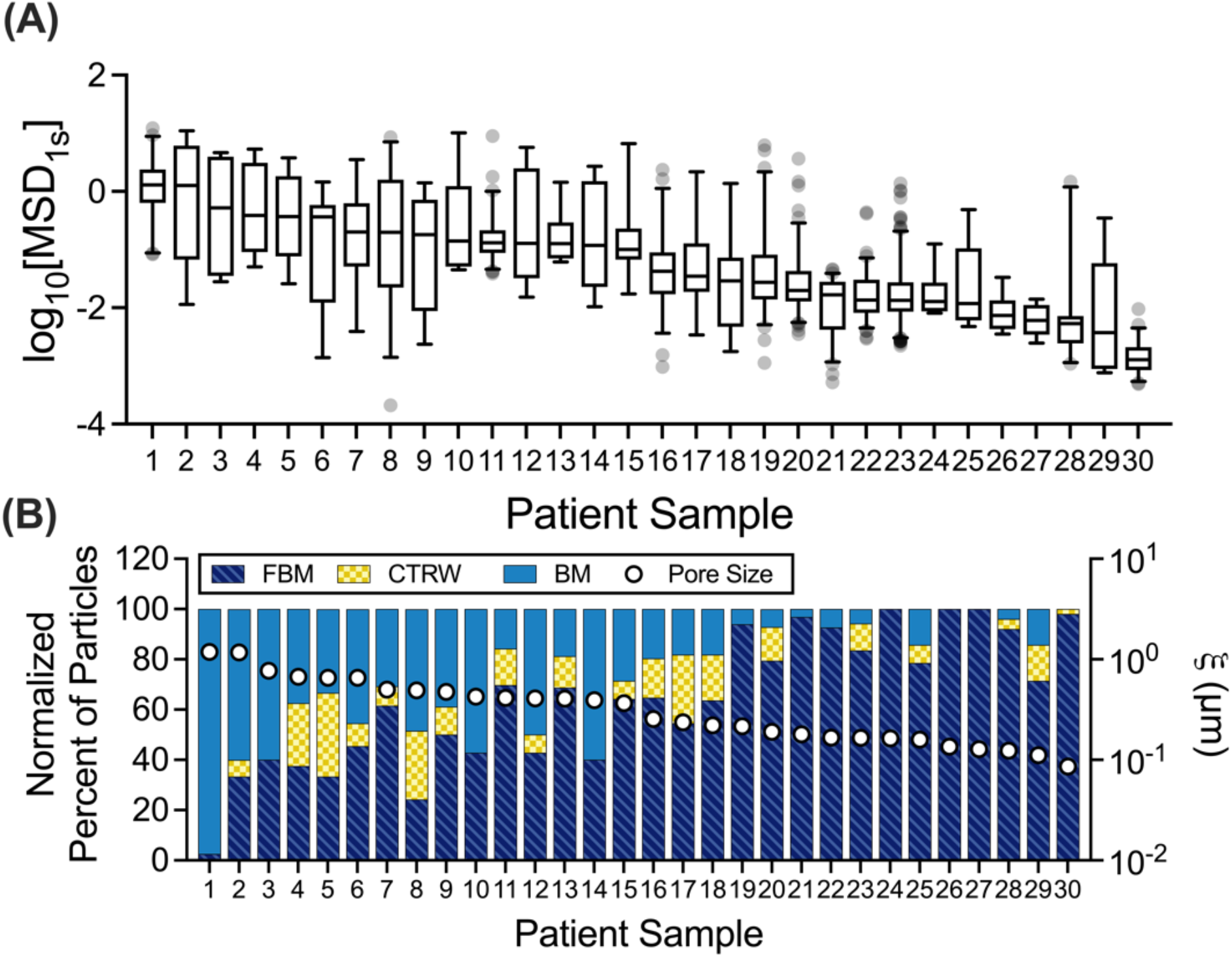
Patient-to-patient variation in NP mobility and diffusion classifications. **(A)** Distribution of log_10_[MSD_1s_] values for NP in each individual patient sample. Patient samples were ordered and numbered by to decreasing median log_10_[MSD_1s_] values. **(B)** Normalized percent of particles exhibiting each type of motion in individual patient samples (left y-axis) compared to the median pore size calculated for each patient sample (right y-axis). Whiskers are drawn down to the 5^th^ percentile, up to the 95^th^ percentile, and outliers are plotted as points.

### Prediction of anomalous diffusion exponents and diffusion coefficients

To further investigate the correlation between diffusion rate of NP in mucus and the classification of motion, we compared log_10_[MSD_1s_] values in each classification (BM, FBM, CTRW) across all samples tested (**Figure 3A**). We observed significant differences in the log_10_[MSD_1s_] values where as expected FBM NP have the lowest diffusivity, BM NP have the highest diffusivity, and diffusivity of CTRW NP was in an intermediate range (i.e., FBM < CTRW < BM). Using the MoNet analysis, we generated the predicted anomalous diffusion exponent (a) for each NP (Figure 3B), which was subsequently used to calculate the effective diffusion coefficient 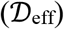 for each NP (Figure 3C). We note α = 1 for all BM NP indicative of normal diffusion and α < 1 for FBM or CTRW NP indicative of anomalous sub-diffusion. Based on measured MSD and α, we find effective diffusion coefficients vary significantly between diffusion types following the same trend as observed in the log_10_[MSD_1s_] values; FBM NP have the lowest effective diffusion coefficients, BM NP have the highest, and the CTRW NP are in an intermediate range.

**Figure 3.**
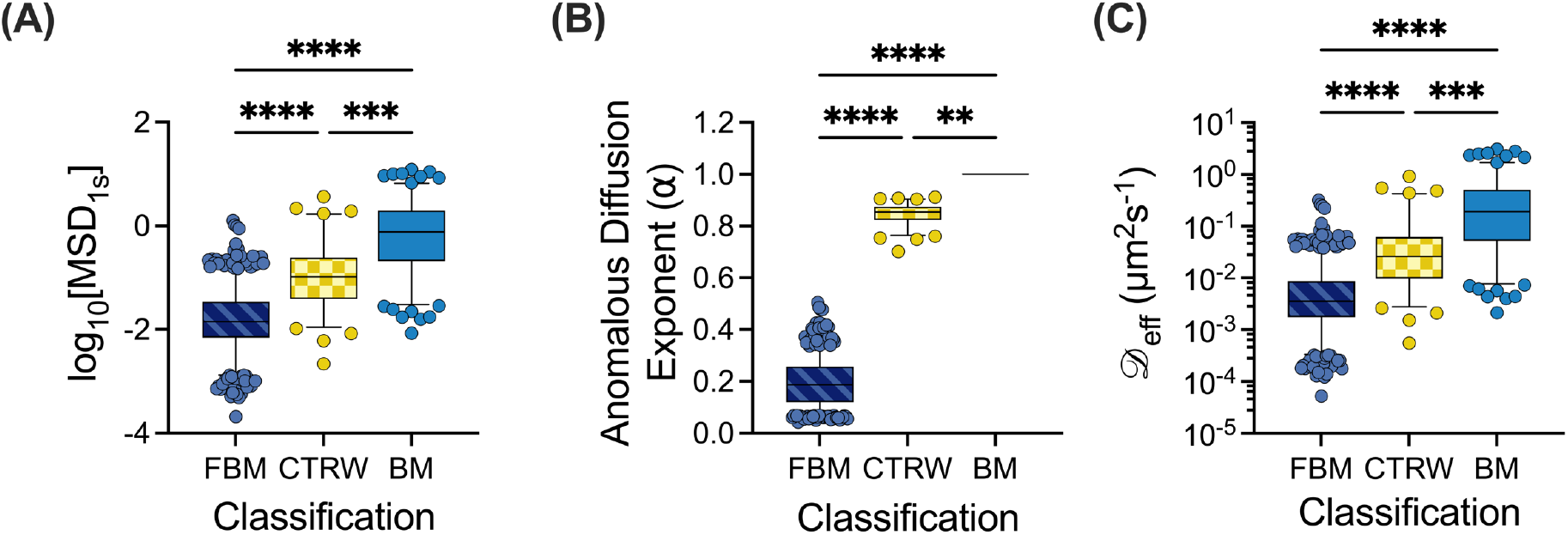
NP mobility in human mucus is highly dependent on the mode of diffusion. **(A)** log_10_[MSD_1s_] values for NP exhibiting each type of motion across all patient samples. **(B-C)** Distribution of predicted anomalous diffusion exponents **(B)** and diffusion coefficients **(C)** for particles exhibiting each type of motion across all patient samples. Whiskers are drawn down to the 5^th^ percentile, up to the 95^th^ percentile, and outliers are plotted as points. Data set statistically analyzed with Kruskal-Wallis test and Dunn’s test for multiple comparisons: *p < 0.05, **p < 0.01, ***p < 0.001, ****p < 0.0001.

### Probability of muco-inert nanoparticles traversing through mucus barrier

Reframing this data in a physiological context, we utilized a mathematical model to predict mean traversal times across the mucus layer developed in previous work (Erickson et al., 2015). The median anomalous diffusion exponents and diffusion coefficients for each type of motion (**Table 1**), were used to generate the survival function (**Figure 4A**), indicating which fraction of particles would remain trapped in 10 μm mucus layer over time. Taking the inverse of the survival function, we obtain the cumulative distribution function (**Figure 4B**), indicating the fraction of particles would cross through the mucus layer over time. Of note, we predict none of the FBM particles would travel across the mucus layer within a 10-hour timespan. We predict all BM NP would reach the underlying cells in ~30 minutes whereas 30–40% of CTRW NP would bypass the mucus layer in 30–60-minute timeframe.

**Figure 4.**
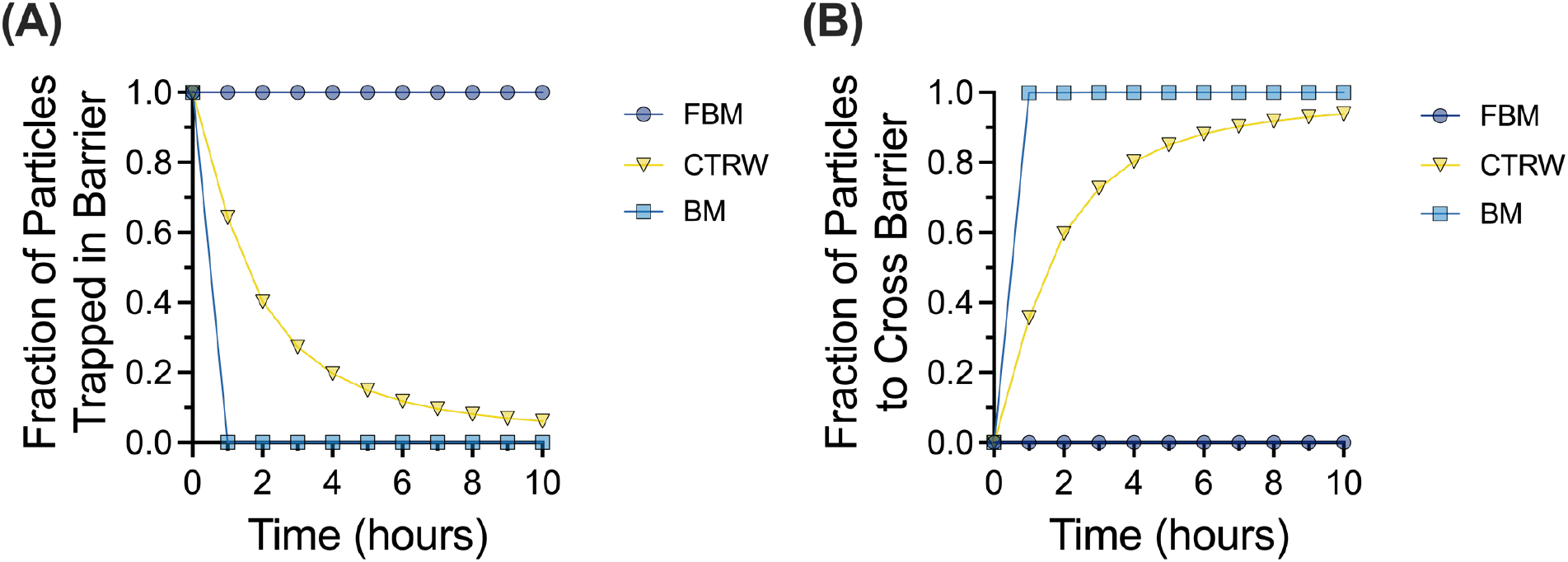
Probability of NP crossing the mucus barrier based on MoNet analysis. **(A)** Survival function showing fraction of particles trapped in mucus barrier layer of 10 μm thickness over time. **(B)** Cumulative distribution function showing the fraction of particles that cross the 10 μm mucus barrier over time.

**Table 1.** Median α and 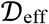 values across all samples.

| | Anomalous Diffusion Exponent ($\alpha$) | Diffusion Coefficient ($\mathcal{D}_{\text{eff}}$, $\mu\text{m}^2\text{s}^{-1}$) |
| --- | --- | --- |
| FBM | 0.1878 | 0.003568 |
| CTRW | 0.8541 | 0.02611 |
| BM | 1 | 0.1925 |

### Variation of percent solids in human mucus effects particle diffusion time

The mucus barrier may also be altered as a function of overall concentration where more concentrated mucus will likely present a more significant barrier to effective nanoparticle drug delivery (Song et al., 2020). To account for the potential changes to the mucus barrier because of varied solids concentrations, 10 samples were grouped based on measured percent solids (**Figure 5A**), with 5% solids as the transition point between low and high percent solids. The log_10_[MSD_1s_] values are grouped by percent solids (**Figure 5B**) and further separated by each type of motion (**Figure 5C**). The α and 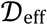 values for each type of motion were calculated for each sample and grouped by the percent solids. The distribution of 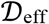 values for NP exhibiting each type of motion is shown for samples grouped by low and high percent solids (**Figure 5D**). Notably, there is a significant difference between the 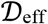 values for FBM NP, with FBM NP in high percent solids samples having lower 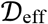 values. The median α and 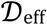 values for low and high percent solids (**Table 2**) were then used in the particle survival analysis. The resulting cumulative distribution function was used with the normalized percentage of particles for each type of motion to calculate the percentage of particles that would be able to cross the 10μm mucus barrier (**Figure 5E**). The resulting percentages indicated that in mucus with low percent solids a larger fraction of BM and CTRW particles are predicted to cross the mucus barrier in a shorter amount of time than in mucus with high percent solids. However, regardless of percent solids, the FBM NP are not predicted to cross the mucus barrier.

**Figure 5.**
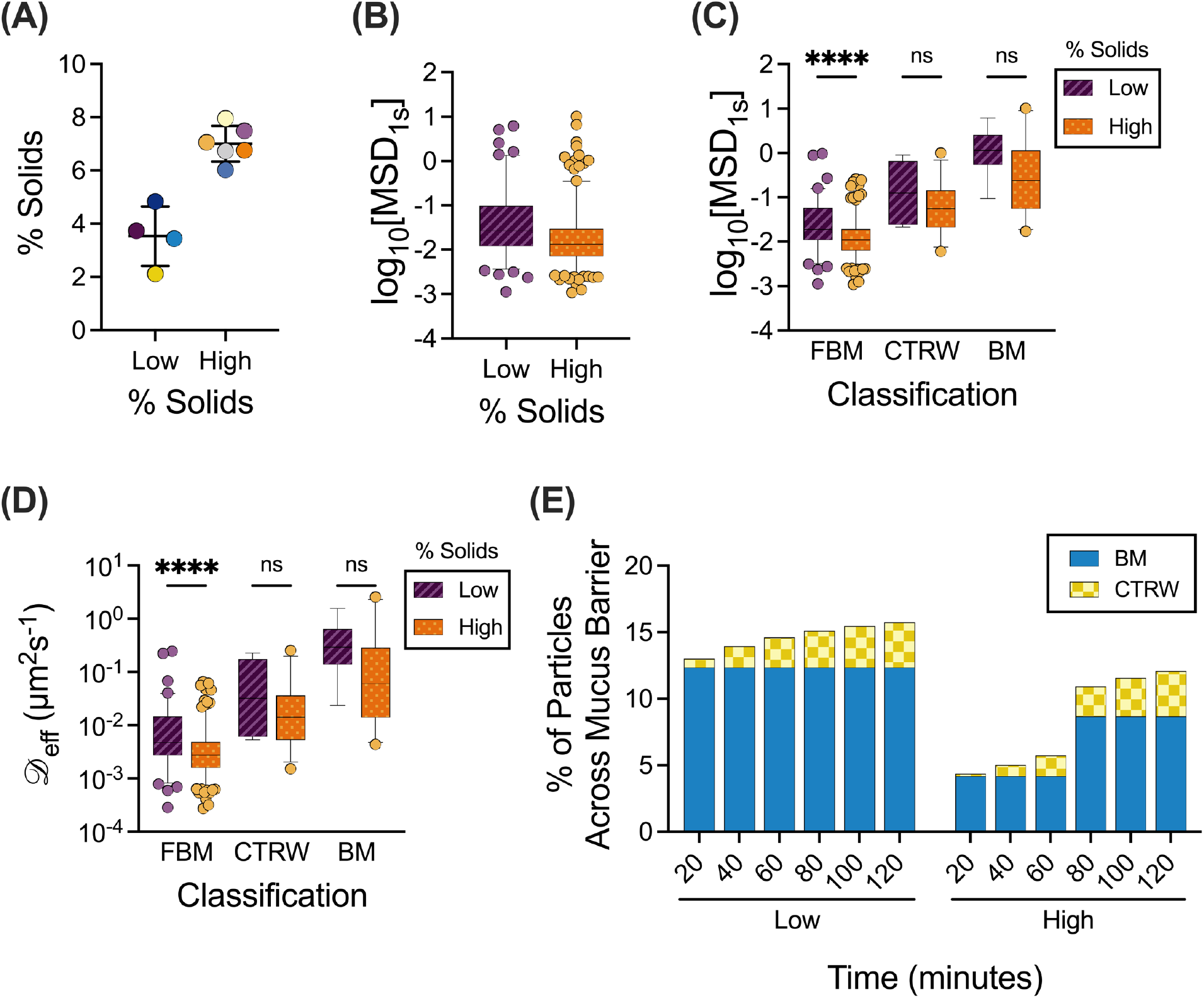
Effect of solids concentration on NP mobility and penetration through the mucus barrier. **(A)** Human mucus samples (n = 10 samples total) classified into low percent solids (n=4 samples) or high percent solids (n=6 samples). Lines indicate mean and standard deviation. **(B-C)** Distribution of log_10_[MSD_1s_] values for NP in patient samples classified as having low or high percent solids **(B)** and NP of classified by motion type and percent solids **(C)**. (**D**) Distribution of effective diffusion coefficients 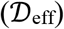 for particles exhibiting each type of motion. **(E)** Percent of particles to cross the 10 μm mucus barrier over time, calculated from the cumulative distribution function and normalized percent of particles. Whiskers are drawn down to the 5^th^ percentile, up to the 95^th^ percentile, and outliers are plotted as points. Data set statistically analyzed with two-tailed Mann-Whitney test: *p < 0.05, **p < 0.01, ***p < 0.001, ****p < 0.0001.

**Table 2.** Median α and 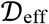 values across samples grouped by percent solids.

| | Anomalous Diffusion Exponent ($\alpha$) | | Diffusion Coefficient ($\mathcal{D}_{\text{eff}}$, $\mu\text{m}^2\text{s}^{-1}$) | |
| --- | --- | --- | --- | --- |
|  | Low % Solids | High % Solids | Low % Solids | High % Solids |
| FBM | 0.2173 | 0.174 | 0.004806 | 0.002777 |
| CTRW | 0.8614 | 0.8641 | 0.03242 | 0.01435 |
| BM | 1 | 1 | 0.2931 | 0.06079 |

### Probability of influenza A virus particles crossing the mucus barrier

We then applied the MoNet and particle survival analyses to a recently published data set for influenza A virus (IAV) particle diffusion in human mucus from 10 different patient donors (Kaler et al., 2022). The trajectories were classified via the MoNet analysis and the percent of the IAV particles exhibiting each type of motion are show in **Figure 6A**. The distribution of 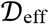 values for the IAV are grouped by motion type and are shown in **Figure 6B**. The median α and 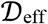 values for the IAV particles (**Table 3**) were then used in the particle survival analysis. The percentage of particles that would be able to cross the 10μm mucus barrier based on the cumulative distribution function and the normalized percent of particles is shown in **Figure 6C**. We found only a small percentage of IAV particles are predicted to cross the mucus barrier, all of which are classified as BM particles.

**Figure 6.**
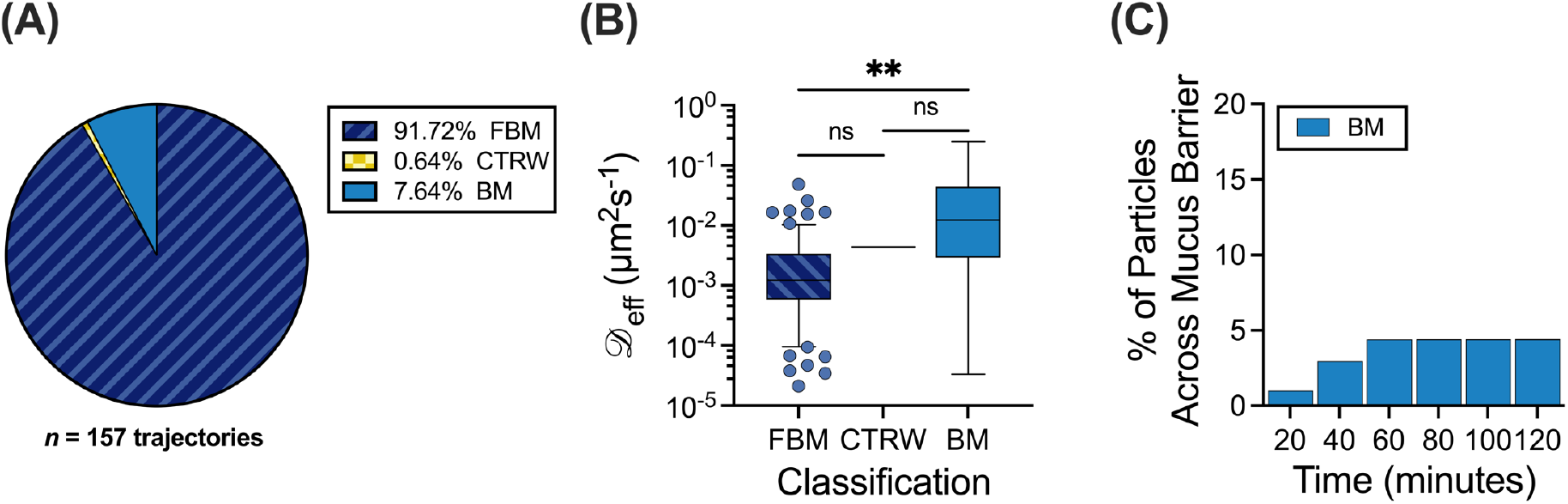
Application to influenza A virus diffusion in human mucus. **(A)** Percent of IAV particles exhibiting each type of motion. **(B)** Distribution of effective diffusion coefficients 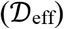 for IAV in human mucus classified by motion type. **(C)** Percent of IAV particles to cross the 10 μm mucus barrier over time, calculated from the cumulative distribution function and normalized percent of particles. Whiskers are drawn down to the 5^th^ percentile, up to the 95^th^ percentile, and outliers are plotted as points. Data set statistically analyzed with Kruskal-Wallis test and Dunn’s test for multiple comparisons: *p < 0.05, **p < 0.01, ***p < 0.001, ****p < 0.0001.

**Table 3.**
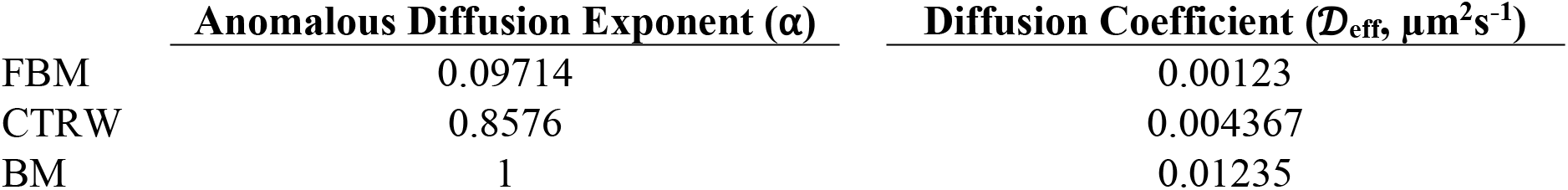
Median α and 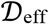 values for IAV in human mucus.

| | Anomalous Diffusion Exponent ($\alpha$) | Diffusion Coefficient ($\mathcal{D}_{\text{eff}}$, $\mu\text{m}^2\text{s}^{-1}$) |
| --- | --- | --- |
| FBM | 0.09714 | 0.00123 |
| CTRW | 0.8576 | 0.004367 |
| BM | 1 | 0.01235 |

## DISCUSSION

In this work, we examined the different types of particle motion that NP exhibited in human airway mucus samples collected and analyzed *ex vivo.* We showed that FBM is the predominant type of motion (**Figure 1**) and identified a potential link between network structure, quantified by pore size, and the predominant type of motion (**Figure 2**). We found clear differences in diffusivity for each type of particle, based on measured log_10_[MSD_1s_] and effective diffusion coefficients (**Figure 3)**. As expected, BM NP traverse the mucus layer most rapidly, whereas CTRW NP can travel through the mucus layer given adequate time (**Figure 4**). However, FBM NP remain trapped in the mucus layer even at longer timescales, up to 3 weeks (data not shown). This suggests that for effective drug delivery, particles would need to exhibit either BM or CTRW movement to traverse the mucus barrier to reach the underlying airway epithelium.

We also found that the percent solids plays an important role in how quickly NP cross the barrier, where a much larger fraction of particles traversed through mucus barrier with sub–5% total solids content (Figure 5). This could be explained by the overall density and reduced mucus network pore size expected in mucus with high percent solids (>5%) which more greatly impedes NP movement. This observation is further corroborated by the increased percentage of CTRW NP and decreased percentage of BM NP in mucus with high percent solids. When considering rapidly moving BM NP, we find the median effective diffusion coefficients for BM NP are ~5-fold higher at low solids concentration as compared to NP dispersed in mucus with high percent solids (**Figure 5D, Table 2**). This is likely due to crowding in mucus with high percent solids as compared to mucus with low percent solids. These findings are also consistent with a previous study by Markovetz et al., in which they found that nanoparticles in human mucus exhibited restricted, sub-diffusive motion, and particle motion restriction increased with increased concentration of solids (Markovetz et al., 2019).

The ramifications of these results are particularly interesting, as previous studies have shown that the percent solids concentration of airway mucus positively correlates with disease severity in obstructive lung diseases such as cystic fibrosis and chronic bronchitis (Boucher, 2019). Thus, one would anticipate nanomedicine used in the treatment of these disorders would encounter a mucus barrier with increased percent solids which could potentially reduce their efficacy However, at very high percent solids, 7-10%, the cilia compresses, causing reduced MCC (Boucher, 2019) and this may provide a larger timeframe for NP to successfully reach the underlying airway epithelial cells. Through these analyses, we can interpret experimental MPT data collected and make more accurate predictions of NP passage accounting for differences in mucus barrier properties in healthy airways and airways of individuals with respiratory disease.

We have also applied these analyses to the diffusion of IAV particles in airway mucus. Intriguingly, while IAV has been shown to be primarily sub-diffusive in previous studies (Yang et al., 2014; Wang et al., 2017), there is a small percentage, 2-4%, of particles that are predicted to cross the mucus barrier in a physiological 30–60-minute time frame, all of which exhibit BM. The outer envelope of IAV contains neuraminidase (NA) and hemagglutinin (HA) which impact their interactions with airway mucus (Vries et al., 2019). While HA binds to sialic acid, which is present on mucins, NA is responsible for cleaving sialic acid. The resulting NA-driven movement likely contributes to the small percentage of particles predicted to cross the mucus barrier (Vries et al., 2019). The remaining particles that are effectively trapped and exhibiting primarily FBM, is likely due to steric and adhesive interactions with the mucus network that NA activity alone is not able to overcome (Kaler et al., 2022). Importantly, our results demonstrate how the diffusion of respiratory viruses in mucus can be analyzed using these methods, allowing for further understanding of infectious disease pathobiology.

In summary, we utilized machine learning and mathematical models to further interpret multiple particle tracking of NP diffusion through the mucus barrier. Comparable to previous reports, we found most NP in mucus exhibit sub-diffusive motion, with the majority exhibiting FBM and these particles are unlikely to reach the airway surface in a physiological timespan. Our results suggest NP exhibiting BM are the primary population expected to penetrate the mucus barrier prior to clearance. However, “hopping” CTRW NP can navigate through the mucus barrier albeit over prolonged time frames. Our results suggest that the fate of 100 nm nanoparticle in the mucus barrier will strongly depend on the percentage of solids and network pore size, which subsequently impacts the diffusion time through the mucus layer. This study establishes a workflow to more rigorously evaluate mucus–penetrating capabilities of NP used in drug delivery applications. We have also shown these analyses can be used in studies of viral trafficking through the mucus barrier to complement standard assays used in infectious disease research.

## METHODS

### Nanoparticle preparation

As previously reported, carboxylate modified fluorescent polystyrene nanoparticles (NP; Life Technologies) with a diameter of 100 nm were coated with 5-kDA methoxy polyethylene glycol (PEG)-amine (Creative PEGWorks) via a carboxyl-amine linkage (Duncan et al., 2016b). Particle size distribution and surface charge was confirmed via dynamic light scattering using the NanoBrook Omni (Brookhaven Instruments). We confirmed dense presence of PEG coating on NP based on the measured zeta potential of 0.04 ± 0.71 mV.

### Human mucus collection

Human mucus was collected under an IRB-approved protocol at the University of Maryland Medical Center (UMMC; Protocol #: HP-00080047). Samples were collected by the endotracheal tube (ET) method, as previously described (Duncan et al., 2016b). ET were collected from 30 donors after intubation as a part of general anesthesia at UMMC. The data presented here are from 6 male and 9 female subjects with mean age of 57 ± 13 years (note: demographic data not available for 15 patients). To collect mucus from ET, the last approximately 10 cm of the tubes were cut, including the balloon cuff, and placed in a 50 mL centrifuge tube. The ET tube was suspended in the tube with a syringe needle and was then spun at 220 g for 30 seconds, yielding 100–300 μL of mucus. Mucus with visible blood contamination was not included in the analysis. Samples were stored in 4°C immediately after collection and imaged within 24 hours.

### Percent solids analysis

To determine the percent solids, 100-150μL of human mucus sample was placed on pre-weighed weigh paper the mass was recorded. The sample was then dried on hotplate for 2 hours or until there was no weight change. The remaining sample was weighed and the percent solids was calculated as the difference between the wet sample and dried sample. Due to collected volume, percent solids were measured for 10 of the 30 human mucus samples.

### Fluorescence video microscopy

Samples were prepared for imaging by placing a vacuum grease coated O-ring on microscope cover glass. The sample was then applied to the center of the well and sealed with a coverslip. For each sample, 1 μL of PEG-coated nanoparticles were added to 20 μL human mucus in the center of the slide well and stirred with a pipette tip prior to imaging. Samples were then equilibrated for 30 minutes at room temperature prior to imaging. Slides were imaged using Zeiss LSM 800 inverted microscope with a 63x water-immersion objective. Multiple 10 second videos were recorded at 33.3 frames per second, for each sample. Similar methods were used for measuring influenza A virus diffusion in human mucus and are described in detail in our previously published work (Kaler et al., 2022).

### Multiple particle tracking (MPT) analysis

Acquired fluorescence microscopy videos were processed using a previously developed MATLAB (The MathWorks, Natick, MA) based analysis code to isolate and track imaged particles (Crocker and Grier, 1996; Schuster et al., 2015; Joyner et al., 2020). For each video, the mean squared displacement (MSD) was calculated as 〈MSD(*τ*)〉 = 〈(*x*^2^ + *y*^2^)〉, for each particle. Due to the nature of MPT, NP were tracked for a maximum of 10 s due to their motion out of the focal plane. To minimize the dynamic and static error in our measurements, a lag time of 1 s was used as a representative value for comparison between conditions. The pore size (*ξ*) can be estimated as 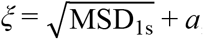, where MSD is the measured MSD at *τ* = 1s and *a* is the NP radius (Song et al., 2021). The same methods were used in the analysis of influenza A virus diffusion in human mucus in our previously published work (Kaler et al., 2022).

### MoNet analysis

Jamali et al. developed a convolutional neural network model named MotionNet (MoNet), which can predict the probability a single trajectory, of length 300 frames, will fall into the diffusion category of Brownian motion (BM), fractional Brownian motion (FBM), or continuous time random walk (CTRW) (Jamali et al., 2021). Once classified by type of diffusion, MoNet can then predict the anomalous diffusion exponent (α) for FBM and CTRW particles (Jamali et al., 2021). The effective diffusion coefficient 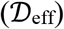 was then calculated for each particle as 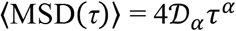, where *α* is the anomalous diffusion exponent predicted by MoNet and τ is the lag time (Tejedor et al., 2010). As some particles move out of frame during imaging, trajectories acquired from the MPT analysis were filtered in MATLAB (The MathWorks, Natick, MA) to isolate trajectories that were tracked for 300 frames. On average, 34 NP trajectories were kept for MoNet analysis per sample tested. In rare cases, a minimum of 5 NP trajectories were kept.

### Particle survival analysis

Erickson et al. developed a model for predicting the first traversal times for nanoparticles using survival functions specific to Brownian motion and fractional subdiffusive motion (i.e. CTRW and FBM) (Erickson et al., 2015). The survival function for diffusive, Brownian motion is given as,

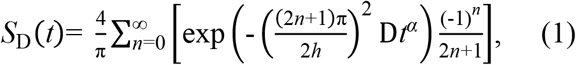

where 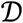 is the diffusion coefficient, α is the anomalous diffusion exponent, and *h* is the absorbing boundary. The survival function for subdiffusive CTRW and FBM is given as,

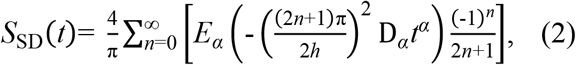

where 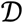 is the diffusion coefficient, α is the anomalous diffusion exponent, *h* is the absorbing boundary, and 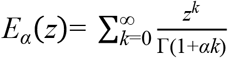 is a Mittag-Leffler function (Erickson et al., 2015). The complement of the survival function, *F*(*t*) = 1 – *S*(*t*), gives the cumulative distribution function (CDF), which is the fraction of particles that cross the absorbing boundary (Erickson et al., 2015). For all the survival functions, a physiological thickness of 10 μm was used for the calculations. The anomalous diffusion exponents (α) and effective diffusion coefficient 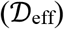 from the MoNet analysis were used to calculate the survival functions for each type of particle using their respective equations. The percentage of particles that would be able to cross the mucus barrier was calculated from the CDF and the normalized percentage of particles for each type of motion using Eq. 3,

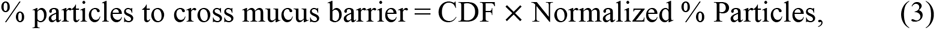

where the CDF and Normalized % of particles are for a single classification of motion.

### Statistical analysis

Data was statistically analyzed using GraphPad Prism 9 (GraphPad Software, San Diego, CA).

### Data availability

Additional data can be made available upon reasonable request.

### Code availability

The particle tracking code used in the current work is available at http://site.physics.georgetown.edu/matlab/. The Python code for MoNet used in this work is available at https://github.com/AliviGitHub/MoNet. The Mathematica code used for the particle survival analysis is available at https://www.cell.com/biophysj/biophysj/supplemental/S0006-3495(15)00546-9.

## AUTHOR CONTRIBUTIONS

L.K., conceived, designed, and performed experiments and analyzed data to determine particle diffusion modes. K.J. conceived, designed, and performed experiments. G.A.D. conceived and designed experiments. L.K. and G.A.D. wrote the article. All authors reviewed and edited the article.

## ACKNOWLEDGEMENTS

This project was funded by NIH R21 AI142050, Burroughs Welcome Fund CASI, and the American Lung Association (to G.A.D.). L.K. was supported by NIH Institutional Training Grant T32 AI089621. K.J. was supported by the Cystic Fibrosis Foundation Postdoctoral Fellowship (JOYNER18FO). We would also like to thank Dr. Irina Timofte and Anu Varghese at UMMC for collecting the human mucus samples used in these studies. The authors have no conflict of interests to disclose.

## Notes

### Competing Interest Statement

The authors have declared no competing interest.

